# HDAC1-3 inhibition reduces CCR2 expression and immunosuppressive function of myeloid-derived suppressor cells

**DOI:** 10.1101/2023.06.01.543360

**Authors:** Zhiqi Xie, Yoshiaki Okada, Daisuke Okuzaki, Naoki Okada, Masashi Tachibana

## Abstract

Myeloid-derived suppressor cells (MDSCs) play a critical role in cancer progression and resistance, making them significant targets for cancer immunotherapy. Although epigenetic regulation by histone deacetylases (HDACs) regulates cell fate and function, the specific roles of HDACs in modulating MDSCs remain poorly understood. We aimed to examine the effects and underlying mechanisms of HDAC on MDSCs using various HDAC inhibitors. HDAC1-3 inhibitors were found to reduce the expression of CCR2, a chemokine receptor that mediates the migration of monocytic (M-)MDSCs to tumors and attenuated the immunosuppressive activity of MDSCs. In an orthotopic hepatocellular carcinoma (HCC) murine model, HDAC1-3 inhibitors reduced the infiltration of M-MDSCs, increased the number of natural killer cells in tumors, and suppressed tumor growth. Our results also suggest that HDAC1-3 inhibitors potentiate the antitumor effects of anti-programmed cell death protein 1 antibodies. To elucidate the molecular mechanisms underlying the inhibition of MDSCs by HDAC1-3 inhibitors, ATAC-seq and RNA-seq analyses were performed. We identified 115 genes that were epigenetically upregulated by HDAC1-3 inhibitors, related to transcriptional regulation and ubiquitination. HDAC1-3 inhibitors further reduced CCR2 protein expression by enhancing ubiquitination-mediated degradation. Our findings reveal a novel mechanism of action of HDAC1-3 inhibitors in MDSCs and suggest a potential combination strategy with immunotherapy for the clinical translation of HCC.

## Introduction

The dysfunction or exhaustion of antitumor immune cells, such as effector T and natural killer (NK) cells, can facilitate immune evasion and tumor progression. Immunotherapy is a promising approach for cancer treatment, particularly immune checkpoint blockade (ICB) therapy, which can reinvigorate antitumor immunity by alleviating the immunosuppression of tumor cells against antitumor immune cells (1). However, most patients still show limited or no response to ICB therapy, probably because it is less effective in relieving immunosuppression mediated by other cells, such as myeloid-derived suppressor cells (MDSCs). MDSCs are a heterogeneous population of myeloid progenitor and immature myeloid cells that are partially blocked in their differentiation and development under tumor conditions, forming two phenotypes: monocytic (M-)MDSCs and polymorphonuclear (PMN-)MDSCs (2). They inhibit antitumor immune responses by secreting regulatory factors, altering immune metabolism, and expressing immune checkpoints, which play crucial roles in the pathogenesis, recurrence, metastasis, immune escape, and drug resistance of multiple cancers (3-5). Studies have revealed that MDSC levels increase in patients with cancer who are resistant to ICB therapy, suggesting that MDSCs promote tumor immune escape via a mechanism distinct from ICB therapy, and that targeting MDSCs could improve the response rate to immunotherapy (6, 7).

Epigenetic regulation plays crucial roles in cancer progression and immune cell function. Histone deacetylases (HDACs) are key molecules that regulate cell fate by removing acetyl groups from lysine residues at the N-termini of histones, modulating gene transcription and expression. Several studies have reported the effects of HDACs on MDSCs. HDACs in mammals are classified into four classes: Zn^2+^-dependent class I (HDAC1, 2, 3, and 8), class IIa (HDAC4, 5, 7, and 9), class IIb (HDAC6 and 10), class IV (HDAC11) HDACs, and NAD^+^-dependent class III (Sirt1-7) HDACs, with distinct structural features (8). HDAC11 deficiency enhances the immunosuppressive activity of MDSCs in tumor-bearing mice (9). Hashimoto *et al*. reported that MS-275, an inhibitor of HDAC1-3, reduced the immunosuppressive activity of PMN-MDSCs, whereas ricolinostat, an inhibitor of HDAC6, reduced the immunosuppressive activity of M-MDSCs in tumor-bearing mice (10). Our previous study revealed that valproic acid (VPA), a class I/IIa HDAC inhibitor (HDACi), significantly reduced the accumulation of M-MDSCs in tumors and attenuated PMN-MDSC immunosuppression (5). Thus, different HDACs play distinct roles in the regulation of MDSCs. However, the effects and mechanisms underlying MDSC regulation by HDAC remain largely unknown. This study aimed to identify the types of HDACs that are important for regulating the accumulation and immunosuppression of MDSCs in tumors.

Hepatocellular carcinoma (HCC) accounts for 75%–85% of liver cancers, with over 80% of patients diagnosed at an advanced stage, and a 5-year survival rate of only 10%–18% (11). The first-line systemic treatment for advanced HCC shifted from multikinase inhibitors, such as sorafenib, to immunotherapy after the Food and Drug Administration approved the combination of a programmed death-ligand 1 monoclonal antibody (atezolizumab) and a VEGF antagonist (bevacizumab) in 2020. However, the objective response rate of immunotherapy is only approximately 30%, which is significantly higher than that of sorafenib (13%); however, 70% of patients with HCC remain unresponsive (12). Therefore, the role and molecular mechanisms of tumor microenvironments in HCC immune evasion needs to be explored. MDSCs contribute to immune evasion and resistance in HCC. For example, MDSC elimination in patients with hepatitis can prevent or delay HCC development (13). Tumor-infiltrating MDSCs inhibit effector T cells, mediate HCC immune escape, and promote tumor recurrence and metastasis (14, 15). MDSC levels were found to increase in patients with HCC resistant to ICB therapy; thus, targeting MDSCs could improve immunotherapy outcomes (16, 17). Therefore, we explored whether strategies targeting HDACs in MDSCs could enhance the efficacy of existing HCC therapies.

## Materials and methods

### Cell lines

The Hepa 1-6 (RCB1638) cell line was obtained from the RIKEN CELL BANK. The cells were thawed and cultured according to the RIKEN CELL BANK guidelines. Dulbecco’s modified Eagle medium (FUJIFILM Wako, Japan) containing 10% fetal bovine serum (FBS) (Gibco, USA) and 1% antibiotic– antimycotic mixed stock solution (100×) (Nacalai Tesque, Japan) was used. The cells were cultured in a CO_2_ incubator at 37°C, saturated vapor pressure, and 5% CO_2_ and used within 1 month of thawing from the early passage (less than three passages in the original vial).

### Experimental animals

The C57BL/6J mice were purchased from Japan SLC (Shizuoka, Japan). They were bred in a specific pathogen-free environment, and 6–8-week-old female mice were selected in the experiments. All animal experiments were conducted in compliance with regulations after obtaining approval from the Osaka University Experimental Animal Committee.

### *In vitro* MDSC differentiation

The bone marrow (BM) cells were differentiated into MDSCs *in vitro*. Briefly, BM cells from C57BL/6J mice were stimulated with 40 ng/mL recombinant GM-CSF (Peprotech, NJ, USA) for 4 days. MS-275 (Carbosynth, UK), TC-H 106 (Tocris, UK), bufexamac (FUJIFILM, Japan), TMP269, PCI-34051, Panobinostat, SAHA, TSA (MedChemExpress, USA), and SIS17 (Selleck, USA) were dissolved in dimethyl sulfoxide (DMSO) (10 mM), diluted with RPMI 1640 medium, and added at the initiation of *in vitro* MDSC differentiation. In examining the proteolytic system, MS-275 or CXD101 was added (250 nM) on day 4 of *in vitro* MDSC differentiation, MG-132 (Selleck) was dissolved in DMSO to a concentration of 100 mM, diluted with RPMI 1640 medium, and added at the same time as the above HDAC inhibitor (5 μM). 3-methyladenine (MA) (Selleck) was freshly adjusted to 20 mM in RPMI 1640 medium in a 50°C water bath and added at the same time as the HDAC inhibitor (1 mM). *In vitro* MDSCs were harvested and after 6 h and subjected to flow cytometry.

### Flow cytometry

After washing various cells with 2% FBS/phosphate buffered saline (PBS), mononuclear cells were treated with TruStain FcX (anti-mouse CD16/32) antibody (Clone 93, BioLegend, USA) and incubated at 4°C for 5 min to block Fc receptors. The following fluorescence-labeled antibodies were then allowed to react at 4°C under light-shielding conditions for 15 min. The cells were then washed twice with 2% FBS/PBS. The 7-amino-actinomycin D Viability Staining Solution (BioLegend) was added to eliminate dead cells, and we analyzed surface antigen expression using a flow cytometer (BD FACSCanto II, BD Biosciences, USA). Data were analyzed using the FlowJo software (BD Biosciences). The following antibodies were used: allophycocyanin (APC)-labeled anti-mouse CD11b (Clone M1/70, eBioscience, USA), Pacific Blue-labeled anti-mouse Gr-1 (Clone RB6-8C5), fluorescein isothiocyanate (FITC)-labeled anti-mouse Ly-6G (Clone 1A8), APC-Cy7-labeled anti-mouse Ly-6C (Clone HK1.4), Pacific Blue-labeled anti-mouse CD45 (Clone 30-F11), FITC-labeled anti-mouse CD8α (Clone 53-6.7), FITC-labeled anti-mouse CD3ε (Clone 145-2C11), APC-labeled anti-mouse NK1.1 (Clone PK136), and phycoerythrin (PE)-labeled anti-mouse CCR2 (Clone SA203G11).

### Analysis of immunosuppressive function of MDSCs

CD4^+^ and CD8^+^ T cells were isolated from the spleen lymphocytes of healthy mice using MojoSort mouse CD4 nanobeads and a MojoSort mouse CD8α selection kit (BioLegend), according to the manufacturer’s protocol. The cells were suspended in RPMI 1640 medium (10% FBS, 2 mM GlutaMAX, 100 units/mL penicillin–streptomycin, 10 mM 4-(2-hydroxyethyl)-1-piperazineethanesulfonic acid, 100 μM nonessential amino acids, 1 mM sodium pyruvate, 55 μM 2-mercaptoethanol; Gibco) and seeded at 1 × 10^5^ cells/well in a 96-well plate. T cell proliferation was stimulated with anti-CD3ε (1 μg/mL)/CD28 (1 μg/mL) antibodies (BioLegend). The anti-CD3ε antibody was coated on each well of the plate via overnight incubation at 4°C with PBS before coculture. CD4^+^ and CD8^+^ T cells were labelled with eFlour670 (eBioscience) and cocultured with *in vitro* MDSCs. The cell proliferation rate was analyzed after 3 days using flow cytometry.

### Establishment of orthotopic HCC mouse model

Hepa 1-6 cancer cells were transplanted into the liver of C57BL/6J mice as previously described (18). Mice were shaved and anesthetized with isoflurane using an experimental animal anesthesia device. They were placed in a supine position on a heat plate, and their abdomens were opened using a median incision. The middle lobe of the liver was exposed to a sterile cotton swab, and 20 μL of Hepa 1-6 cells/matrigel solution (Corning) (5 × 10^7^ cells/mL) was injected using a microsyringe (100 μL; Agilent, Japan). The injection site was pressed with a sterile cotton swab to stop bleeding and the middle lobe was returned to its original position. Finally, the abdomen was closed by continuous suturing of the abdominal wall with curved suture needles (Natsume Seisakusho), and the epidermis was fixed with an experimental animal automatic suture device (using forceps and suture clips of 9 mm, Natsume Seisakusho). Ten days after the inoculation of Hepa 1-6 cells, the vehicle solution or MS-275 (5 mg/kg; Selleck) and CXD101 (5 mg/kg; MedChemExpress) were intraperitoneally administered daily for 10 days. MS-275 and CXD101 were prepared by dissolving them in 1% DMSO, 15% PEG300, and PBS. Anti-PD-1 antibody (Clone: J43) or IgG antibody (200 μg/mouse, Bio X Cell) was intraperitoneally administered every 3 days from 10 days after the inoculation of Hepa 1-6 cells. On day 21 after Hepa 1-6 transplantation, the livers were removed and photographed. The tumor was excised, and its weight was measured.

### Mononuclear cell isolation

Spleen: mice spleens were collected and the cells were dispersed on a 70-μm cell strainer. After centrifugation at 330 × g, 4°C for 5 min, the supernatant was aspirated. The obtained cells were resuspended in ammonium–chloride–potassium (ACK) lysis buffer to remove red blood cells and washed with 2% FBS/PBS to obtain splenic mononuclear cells.

Tumor: mice tumors were finely minced using scissors and dispersed in 100 units/mL collagenase I (Funakoshi, Japan), 2% FBS/PBS, and incubated at 37°C for 30 min for cell dispersion. The tissue fragments were removed using a 70-μm cell strainer and after centrifugation at 330 × g, 4°C for 6 min, the supernatant was aspirated. The cells were suspended in ACK lysis buffer to remove red blood cells and washed with 2% FBS/PBS to obtain mononuclear tumor cells.

Peripheral blood: peripheral blood collected from mice fundi was suspended in ACK lysis buffer to remove red blood cells and after centrifugation at 400 × g, 4°C for 5 min, the supernatant was aspirated. This procedure was repeated once to obtain peripheral blood mononuclear cells.

### ATAC-seq analyses

M-MDSCs (CD11b^+^Ly-6C^hi^Ly-6G^-^; purity > 95%) were sorted from isolated tumors using JSAN (KS-Techno, Japan). ATAC-seq libraries were prepared using an ATAC-seq Kit (Active Motif, USA), according to the manufacturer’s protocol. Briefly, 1 × 10^5^ M-MDSCs were lysed with ATAC lysis buffer and nuclei were centrifuged at 500 × g, 4°C for 10 min. Nuclei were then suspended in 50 μL of Tagmentation Master Mix and incubated at 37°C for 30 min with shaking. DNA was purified using a DNA purification column and polymerase chain reaction (PCR) was performed for 10 cycles to introduce sequencing adapters into the open chromatin regions. The library DNA was purified using solid-phase reversible immobilization beads. Molecular weights were checked, and libraries were quantified using a Bioanalyzer and submitted for next-generation sequencing (NGS) to the Genome Information Research Center, Research Institute for Microbial Diseases, Osaka University. FASTQ files were received and ATAC-seq data were analyzed using Active Motif’s Basepair Portal web tool with an automated pipeline. The ATAC-seq libraries had an average of > 70 million reads per sample. Sequenced reads were mapped to the mouse reference genome mm10 using Bowtie2, and differential analyses and gene annotation were performed using MACS2 (default options: no model; *p* < 0.00001; shift: 0). In this analysis, promoters were defined as those located 5 kb upstream of the transcription start site. ATAC-seq tracks were visualized using the Integrative Genomics Viewer (v.2.11.0).

### RNA-seq analyses

Total RNA was extracted from M-MDSCs using the miRNeasy Mini Kit (Qiagen, Germany). RNA libraries were prepared using TruSeq stranded mRNA sample prep kit (Illumina, USA) and sequenced on Illumina NovaSeq 6000 platform (101 bp single-end mode). RNA extraction for NGS was outsourced to the Genome Information Research Center of the Research Institute for Microbial Diseases, Osaka University, Japan. RNA-seq data were analyzed using the BioJupies web tool and a volcano plot was generated. Genes with adjusted *p* < 0.05 and fold change > 1.5 were extracted as significantly differentially expressed genes.

### Quantitative real-time (qRT)-PCR

Total RNA was extracted from MDSCs using TRIzol reagent (Thermo Fisher Scientific, USA), and reverse transcription was performed using the QuantiTect Reverse Transcription Kit (QIAGEN, Germany). The cDNA of each gene was amplified using TB Green Premix Ex Taq II (TaKaRa, Japan), and PCR was performed using the CFX96 Real-Time PCR system (Bio-Rad, USA). Expressions were compared using the ΔΔCt method and mRNA expressions were normalized to those of mouse *Gapdh* mRNA. The primers used are listed as follows: *Gapdh*, 5′-TGACCTCAACTACATGGTCTACA-3′ (forward); 5′-CCGTGAGTGGAGTCATACTGG-3′ (reverse). *Ccr2*, 5′-ATCCACGGCATACTATCAACATC-3′ (forward); 5′-CAAGGCTCACCATCATCGTAG-3′ (reverse).

### Statistical analyses

GraphPad Prism (GraphPad Software, USA) was used for statistical analyses and *p*-values were calculated using Student’s *t* test, one-way ANOVA, two-way ANOVA, log-rank test, or Pearson’s correlation coefficient test.

## Results

### HDAC1-3 inhibitors reduced CCR2-expressing M-MDSCs

The expression of CCR2, a chemokine receptor on the surface of M-MDSCs, regulates their migration toward tumors and promotes tumor progression. We have previously reported that VPA can significantly reduce CCR2-expressing M-MDSCs, specifically by reducing their accumulation in tumors and inhibiting the growth of lymphoma and melanoma. Therefore, we conducted further studies to investigate HDAC inhibitors that are important for CCR2 expression on MDSCs. Consistent with previous studies, CCR2 was absent in PMN-MDSCs but was present in M-MDSCs (Fig. 1A). VPA treatment significantly reduced the proportion of CCR2^+^ M-MDSCs (Fig. 1B). HDAC1-3 inhibitors (MS-275, CXD101, and TC-H 106) and pan-HDAC inhibitors reduced the CCR2^+^ population in M-MDSCs, whereas the other HDAC inhibitors exhibited no effects (Fig. 1B). MS-275, CXD101, and TC-H 106 downregulated the expression of CCR2 at low concentrations without affecting cell proliferation in a dose-dependent manner (Fig. 1C, S1). Thus, HDAC1-3 enhanced CCR2 expression on MDSCs.

**Fig. 1.**
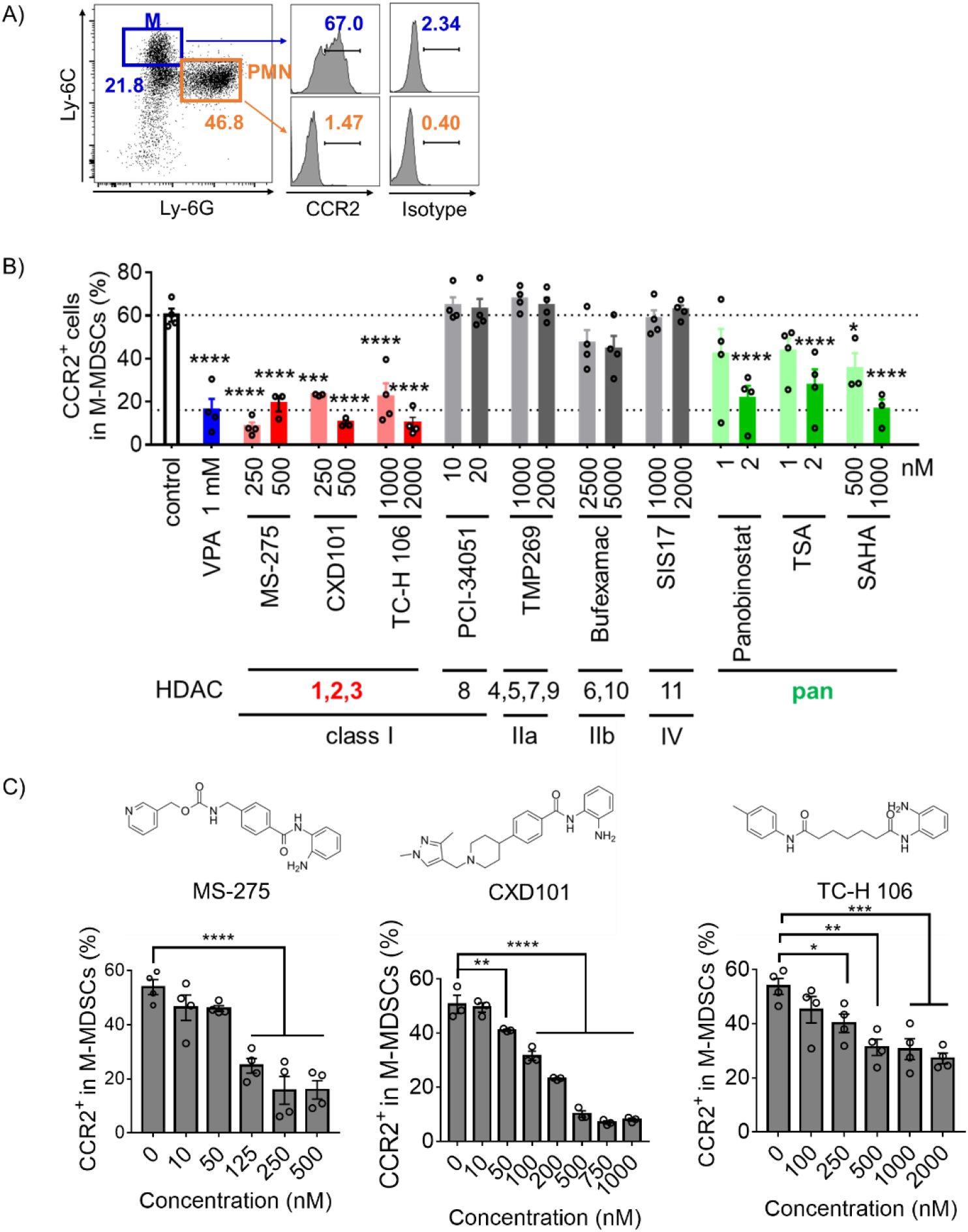
HDAC1-3 inhibitors downregulate CCR2 expression in M-MDSCs. After 4 days of culture in medium supplemented with GM-CSF (40 ng/mL) with or without the addition of different concentrations of HDAC inhibitors, CCR2 expression on M-MDSCs was assessed using flow cytometry. **A)** Gating strategy used for analysis of CCR2 on M-MDSCs and PMN-MDSCs. **B)** CCR2 expression in M-MDSCs treated with HDAC inhibitors. Data represent mean ± S.E.M., pooled from three or four independent experiments (\*\*\**p* < 0.001, \*\*\*\**p* < 0.0001 by one-way ANOVA, compared with control group). **C)** Molecular structures of HDAC1, 2, 3 inhibitors (MS-275, CXD101, and TC-H 106) are shown above. CCR2 expression on M-MDSCs treated with different concentrations of the HDAC inhibitors was assessed. Data represent mean ± S.E.M., pooled from two independent experiments (\**p* < 0.05, \*\**p* < 0.01, \*\*\**p* < 0.001, \*\*\*\**p* < 0.0001 by one-way ANOVA, compared with the group without additional HDAC inhibitor).

### HDAC1-3 inhibitors attenuated the immunosuppressive function of MDSCs

We examined the HDAC inhibitors that are important for regulating the immunosuppressive function of MDSCs. As previously reported, control MDSCs without HDAC inhibitors potently inhibited T cell proliferation, whereas VPA-treated MDSCs exhibited an impaired suppressive function (Fig. 2A, B). MDSCs that were stimulated with HDAC1-3 inhibitors but not with HDAC8 inhibitors enhanced T cell proliferation compared to control MDSCs, as did VPA (Fig. 2A, B). MDSCs stimulated with the class IIb inhibitor bufexamac also exhibited increased T cell proliferation compared to control MDSCs. Conversely, MDSCs treated with the class IIa HDAC inhibitor (TMP269-treated MDSCs) suppressed T cell proliferation more than control MDSCs (Fig. 2A, B), indicating that the immunosuppressive function of MDSCs may be increased by class IIa HDAC inhibition. SIS17-treated MDSCs (a class IV HDAC inhibitor) exhibited a stronger suppression of CD8^+^ T cell proliferation than control MDSCs (Fig. 2A). Thus, different HDACs have different effects on the immunosuppressive functions of MDSCs, particularly targeting HDAC1-3 or class IIb HDACs can inhibit MDSC-suppressive activity. Therefore, HDAC1-3 inhibitors decreased CCR2 expression on MDSCs, attenuated their immunosuppressive function, and were considered effective MDSC inhibitors.

**Fig. 2.**
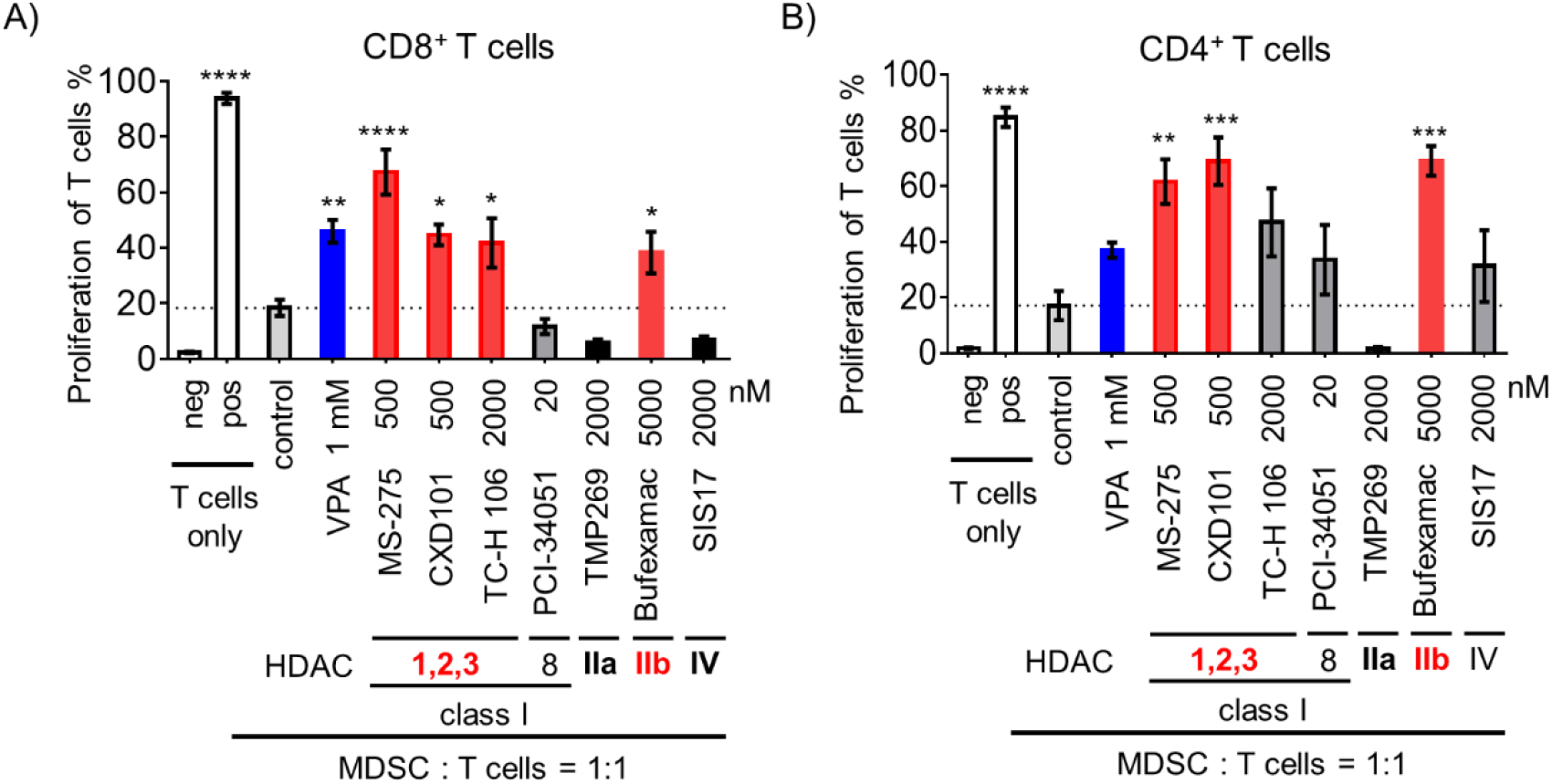
The effect of HDAC inhibitors on the immunosuppressive activity of MDSCs. **A, B)** MDSCs treated with different HDAC inhibitors were combined in a 1:1 ratio with eFluor 670-labeled CD4^+^ or CD8^+^ T cells, followed by stimulation with anti-CD3ε/anti-CD28 antibodies. Data represent mean ± S.E.M. of three independent experiments (one-way ANOVA: \**p* < 0.05, \*\**p* < 0.01, \*\*\**p* < 0.001, and \*\*\*\**p* < 0.0001, compared with control group). Neg, negative control (T cell only, without anti-CD3ε/anti-CD28 antibodies stimulation); pos, positive control (T cell only, with anti-CD3ε/anti-CD28 antibodies stimulation).

### Antitumor effects of HDAC1-3 inhibitors in Hepa 1-6 HCC mouse model

To investigate the potential of HDAC1-3 inhibitions in cancer therapy, we analyzed the expression and prognostic value of HDAC1, 2, and 3 in various cancers using the GEPIA2 web tool based on The Cancer Genome Atlas database (19). We found that only in liver HCC (LIHC), HDAC1, 2, and 3 expression was significantly associated with survival rate and was higher in tumor tissue than normal tissue (Fig. S2A-G). Using TIMER 2.0 (20), we observed a positive correlation between HDAC1, 2, and 3 expression and tumor-infiltrating MDSCs in LIHC (Fig. S2H). These findings suggest that high levels of MDSCs expressing HDAC1, 2, and 3 in LIHC may exacerbate disease outcomes.

To further investigate the antitumor effects of the HDAC1-3 inhibitors, MS-275 and CXD101, Hepa 1-6 orthotopic HCC model mice were used (Fig. 3A). MS-275 and CXD101 were intraperitoneally administered daily for 10 days to Hepa 1-6 transplanted mice. They were found to inhibit tumor growth compared to that in the control mice (Fig. 3B). These inhibitors decreased CCR2 expression on M-MDSCs from the spleen, blood, and tumors. MS-275 tended to reduce M-MDSCs in the blood and tumors, and CXD101 treatment significantly reduced these numbers (Fig. 3C-D). However, PMN-MDSCs from each tissue did not exhibit any differences (Fig. S3A). HDAC1-3 inhibitors did not increase CD8^+^ T cells in tumors but significantly increased NK cells (Fig. 3E, S3B). Furthermore, the number of M-MDSCs was positively correlated with tumor weight, indicating that M-MDSCs are key cells in HCC progression (Fig. 3F). *In vitro* treatment of Hepa 1-6 cells with MS275 or CXD101 resulted in inhibition of their proliferation in a dose-dependent manner (Fig. S3C). However, even at low concentrations, they had little effect on cancer cells (Fig. S3C) and strongly affected MDSCs (Fig. 1C). Therefore, the anticancer effects of MS275 and CXD101 *in vivo* may be mediated by MDSCs rather than by cancer cells. Thus, HDAC1-3 inhibitors may reduce the number of M-MDSCs and relieve the tumor immune suppression microenvironment, thereby limiting HCC progression.

**Fig. 3.**
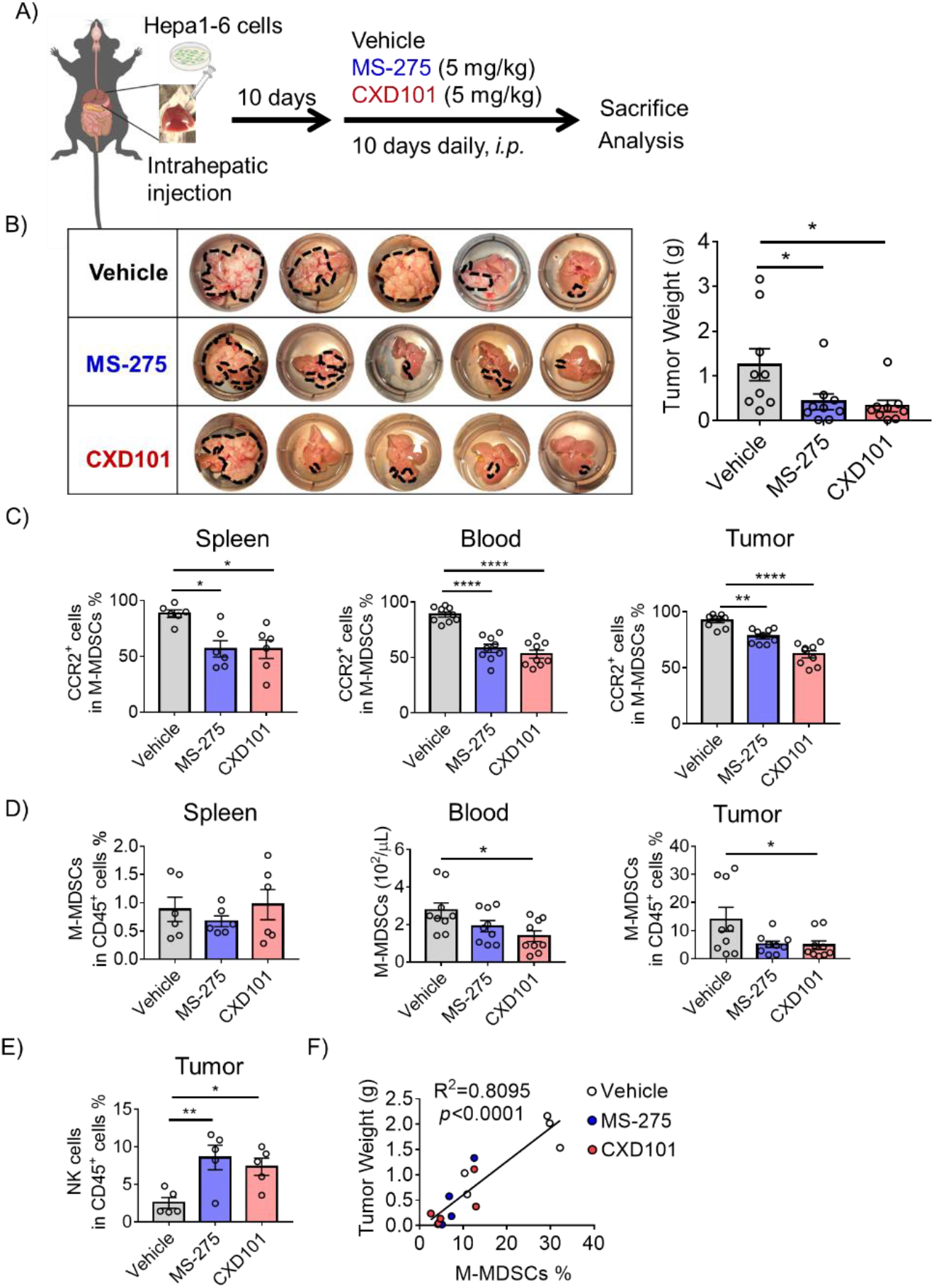
HDAC1-3 inhibitors impaired Hepa 1-6 tumor development. **A)** Orthotopic mouse model using Hepa 1-6 cells was established and administered drugs as per the dosing schedule. **B)** Endpoint tumor weight was measured with representative images displaying tumor morphology. Data are mean ± S.E.M., pooled from two independent experiments with n = 9 per group (\**p* < 0.05 by one-way ANOVA). **C)** The proportion of CCR2^+^ cells among total M-MDSCs from spleen, blood, and tumors were assessed using flow cytometry. Data represent mean ± S.E.M. of one (spleen) or two (blood, tumor) independent experiments (\**p* < 0.05, \*\**p* < 0.01, \*\*\*\**p* < 0.0001 by one-way ANOVA). **D)** Flow cytometry of the M-MDSCs (CD11b^+^Ly-6G^-^Ly-6C^hi^) in spleen, blood, and tumors. Data represent mean ± S.E.M. of one (spleen) or two (blood, tumor) independent experiments (\**p* < 0.05 by one-way ANOVA). **E)** Flow cytometry of the tumor NK cells (CD3ε-NK1.1^+^) in CD45^+^ live cells. Data represent mean ± S.E.M. of two independent experiments (\**p* < 0.05, \*\**p* < 0.01 by one-way ANOVA). **F)** Correlation between M-MDSCs and NK cells in Hepa 1-6 tumor sites was determined using Pearson’s correlation coefficient test. **G)** Correlation between M-MDSCs and tumor weight in Hepa 1-6 tumor sites was determined using Pearson’s correlation coefficient test.

Whether HDAC1-3 inhibitors could enhance the antitumor effects of anti-PD-1 antibodies was examined. CXD101 alone or anti-PD-1 antibody alone also inhibited tumor progression, but the tumors almost completely regressed in the combination group (Fig. 4). Therefore, HDAC1-3 inhibitors potentiate the antitumor effects of anti-PD-1 antibodies in an orthotopic HCC mouse model.

**Fig. 4.**
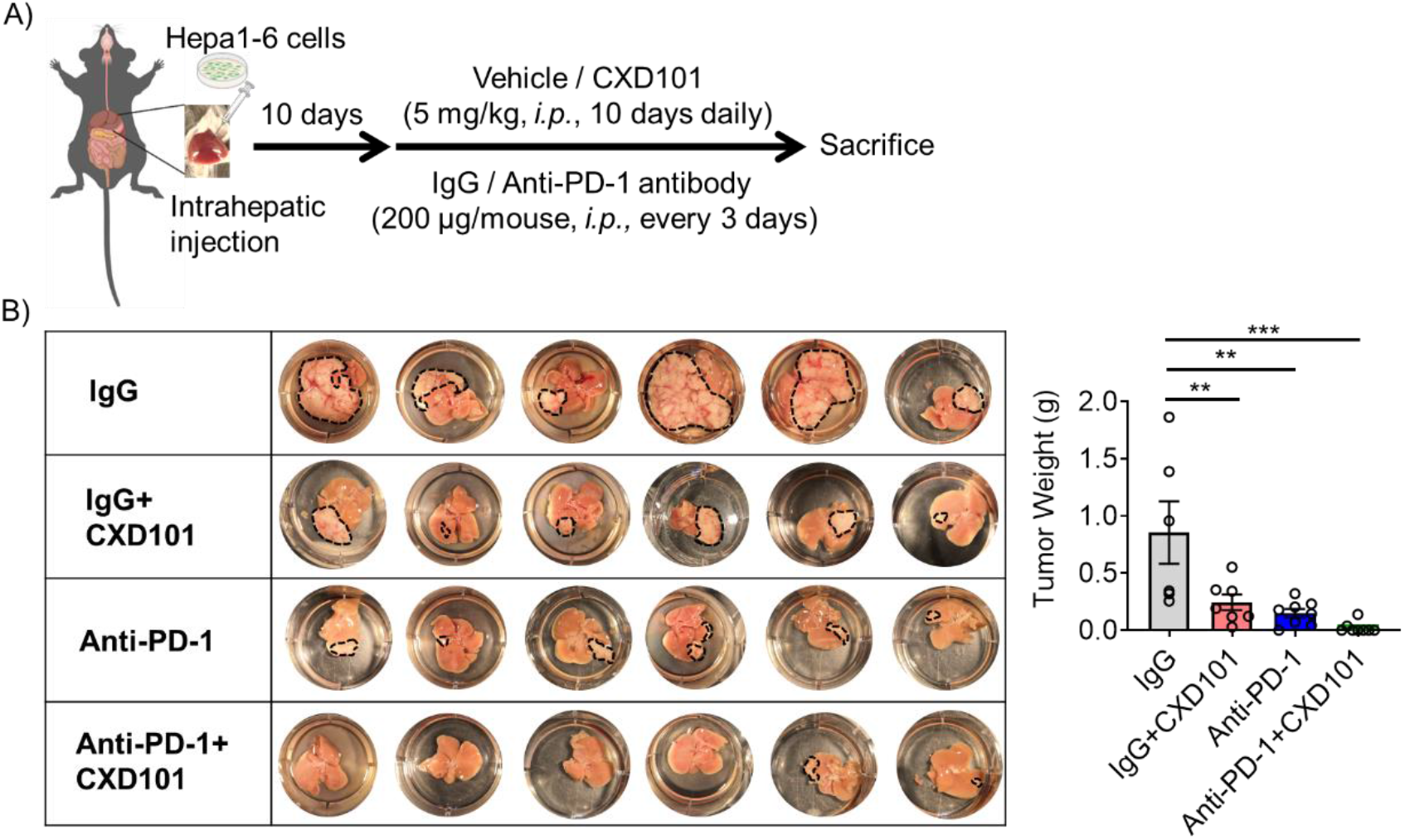
HDAC1-3 inhibitors enhance the antitumor effect of anti-PD-1 immunotherapy. **A)** Ten days after inoculation with Hepa 1-6 cells, the mice were administered intraperitoneal injection of vehicle, MS-275, or CXD101 (5 mg/kg) daily in combination with an anti-PD-1 antibody or IgG (200 μg/mouse) every 3 days until day 20. **B)** Endpoint tumor weight was measured with representative images displaying tumor morphology. Data represent mean ± S.E.M., pooled from two independent experiments with n = 6–8 per group (\*\**p* < 0.01, \*\*\**p* < 0.001 by two-way ANOVA).

### ATAC-seq and RNA-seq analyses of M-MDSCs treated with HDAC1-3 inhibitors

To further elucidate the molecular mechanisms underlying HDAC1-3 inhibitors by MDSCs, we performed ATAC-seq and RNA-seq analyses of M-MDSCs isolated from Hepa 1-6 tumor-bearing mice treated with MS-275 or CXD101 (Fig. 5A). ATAC-seq identifies open chromatin regions using fragmentation and sequencing. ATAC-seq peaks were detected as open chromatin in regions where histones were acetylated by HDAC inhibition (Fig. 5B). In the MS-275-treated group, there were 19,195 peaks, of which 5,387 (28.1%) were in the promoter region (Fig. 5C). In the CXD101-treated group, there were 6,389 peaks, and 1,631 (25.5%) were in the promoter region (Fig. 5D). RNA-seq was conducted to investigate changes in HDAC expression following MS-275 or CXD101 treatment and to identify genes with open chromatin promoters and increased expression. MS-275 treatment resulted in 1716 upregulated genes compared to those in the vehicle group (Fig. 5E). Intersection with the ATAC-seq analysis yielded 775 common genes (Fig. 5F). CXD101 treatment resulted in the upregulation of 832 genes (Fig. 5G), with ATAC-seq peaks observed in the promoter regions of 125 genes (Fig. 5H). Among these, 115 genes were common to both the MS-275- and CXD101-treated groups and were considered candidates for epigenetic regulation by HDAC1-3 in M-MDSCs (Fig. 5I).

**Fig. 5.**
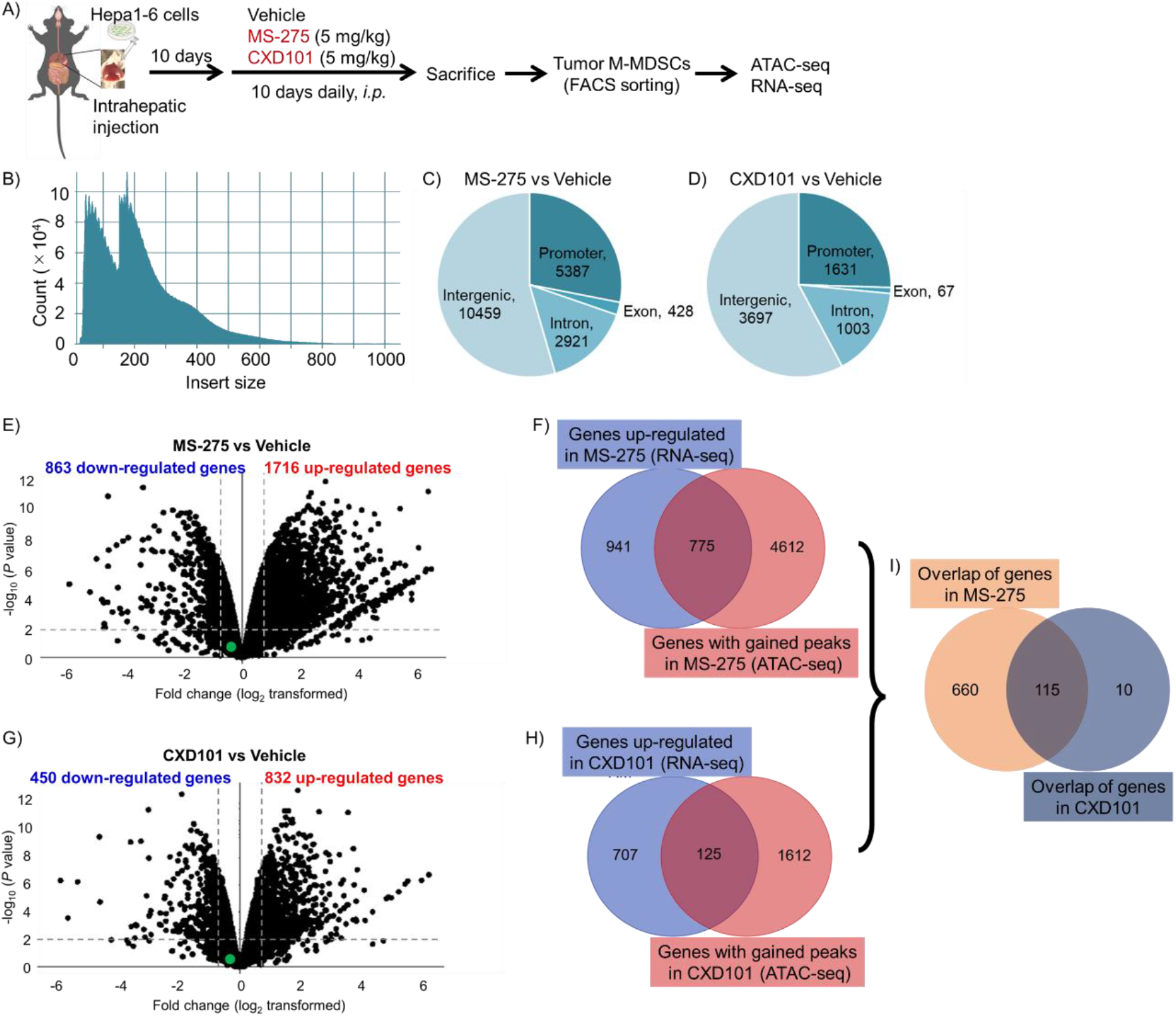
Identifying HDAC1-3 inhibitors-induced M-MDSC epigenetic changes. **A)** The drug dosing schedule is shown. M-MDSCs sorted from Hepa 1-6 tumor site were used for ATAC-seq and RNA-seq. **B)** Insert size distributions of ATAC-seq data indicating clear nucleosome phasing. **C-D)** Proportions and number of the ATAC-seq peaks annotated to different genomic regions in MS-275 and CXD101 groups versus the vehicle group (n = 2 biological replicates). Green dot represents *Ccr2*. **E, G)** Volcano plots showing differences in mRNA expression profiles of MDSCs (n = 2 biological replicates). **F)** Venn diagram showing an overlap of genes with significantly gained ATAC-seq signal in the MS-275 group and upregulated genes in the MS-275 group (RNA-seq) compared with the vehicle group. **H)** Venn diagram showing overlap of genes with significantly gained ATAC-seq signal in the CXD101 group and upregulated genes in the CXD101 group (RNA-seq) compared with the vehicle group. **I)** Venn diagram showing genes overlapping in both MS-275 and CXD101 groups.

Gene ontology enrichment analysis (21) of the 115 identified genes revealed annotations related to transcriptional regulation and ubiquitination (Fig. 6A). The ubiquitination system controls >80% of protein degradation. We first quantified *Ccr2* mRNA expression in M-MDSCs and found that HDAC1-3 inhibitors did not significantly alter *Ccr2* mRNA expression (Fig. 6B, C), suggesting that HDAC1-3 inhibitors did not affect *Ccr2* transcriptional activity. We used the proteasome inhibitor MG-132 and revealed that MG-132 prevented the CXD101-induced decrease in CCR2 expression in M-MDSCs. However, the autophagy inhibitor 3-MA did not prevent the HDAC1-3 inhibitor-induced decrease in CCR2 expression. These results revealed that HDAC1-3 reduced CCR2 protein levels by degrading it via the ubiquitin-proteasome system.

**Fig. 6.**
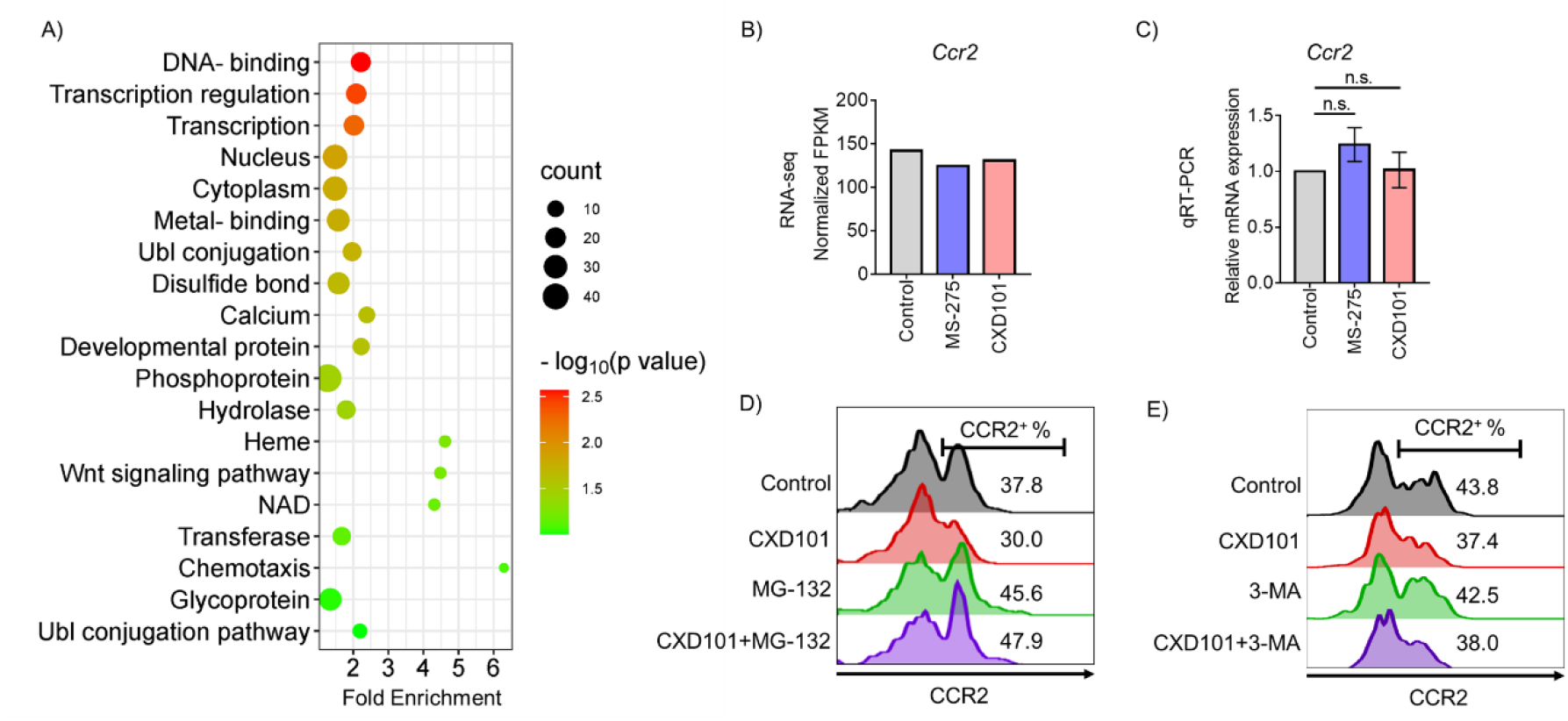
Proteasome-dependent degradation regulates CCR2 expression in M-MDSCs. **A)** DAVID analyses of the identified 115 genes using the UP_KEYWORDS Functional Category (count > 2 with *p* < 0.05). **B)** Normalized FPKM of *Ccr2* gene from RNA-seq (n = 2 biological replicates). **C)** Total RNA was extracted from *in vitro* M-MDSCs, and qRT-PCR was performed to measure the mRNA expression of *Ccr2* (n = 4 biological replicates). **D)** Flow cytometry of CCR2 expression on *in vitro* M-MDSCs after 6 h of treatment with CXD101 (250 nM), MG-132 (5 μM), or CXD101 and MG-132. **E)** Flow cytometry of CCR2 expression on *in vitro* M-MDSCs after 6 h of treatment with CXD101 (250 nM), 3-MA (1 mM), or CXD101 and 3-MA.

## Discussion

MDSCs have been implicated in cancer progression and conferring resistance to chemotherapy, radiotherapy, and ICB therapy. Therefore, the targeting of MDSCs is a significant challenge in cancer immunotherapy. In this study, we examined the epigenetic control mechanisms underlying cell fate and function, and elucidated the role of HDACs in regulating MDSCs. We found that HDAC1-3 inhibition leads to the degradation of CCR2 in M-MDSCs via the ubiquitin-proteasome system. This suggests that the HDAC1-3 inhibitors, MS-275 and CXD101, downregulate CCR2 expression in M-MDSCs and decrease M-MDSCs in tumors. Consistent with our results, MS-275 administration to patients with breast cancer resulted in a greater reduction in M-MDSCs than in PMN-MDSCs (22). Similar to other cancers (23, 24), VPA or HDAC1-3 inhibitors alone exhibited antitumor effects, which were further enhanced when combined with ICB therapy in HCC. Thus, targeting MDSCs and inhibiting HDAC1-3 are promising strategies for cancer immunotherapy.

CCR2 plays a crucial role in regulating M-MDSC function in cancer by affecting tumor growth, metastasis, and immunotherapy resistance (25, 26). Despite its importance, the molecular mechanisms underlying CCR2 expression remain poorly understood. Study has revealed that FOXO1 directly binds to the CCR2 promoter and activates its transcription (27). Our findings reveal a novel regulatory mechanism by which the HDAC inhibitors MS-275 and CXD101 do not affect *Ccr2* transcription but instead reduce CCR2 protein levels by enhancing its ubiquitination-mediated degradation. Although further investigation is necessary to identify the E3 ligase of CCR2 and elucidate the mechanisms by which HDACs modulate E3 ligase activity in CCR2 regulation, our results provide novel insights into the intricacies of CCR2 regulation and suggest that HDAC inhibitors hold therapeutic promise for M-MDSC targeting in cancer.

Our results demonstrated that the inhibition of HDAC1-3 or class IIb HDACs attenuated the immunosuppressive function of MDSCs. Considering VPA lacks class IIb HDAC inhibitory activity, we hypothesized that HDAC1-3 inhibition would reduce MDSC immunosuppression. Additionally, HDAC6 is highly expressed in M-MDSCs and selective inhibitors decrease their immunosuppressive function in tumor-bearing mice (10). This is consistent with our finding that class IIb HDAC inhibitors, including HDAC6, reduce MDSC immunosuppression. However, the mechanisms through which HDACs regulate the immunosuppressive function of MDSCs remain unclear and require further investigation. Additionally, the differential effects of HDACs on MDSC subtypes and their underlying mechanisms should be explored. In conclusion, our evidence suggests that HDAC1-3 inhibitors decrease CCR2 expression in MDSCs, attenuate their immunosuppressive function, and are effective MDSC inhibitors.

HCC is a significant global health issue, ranking fifth in cancer incidence and third in mortality rates. Despite recent targeted and immunotherapeutic approaches, HCC remains a highly refractory and recurrent disease requiring more effective treatment. MDSCs are potential targets for HCC therapy; they are immunosuppressive cells accumulating in the tumor microenvironment and are negatively correlated with treatment response and survival. We found that HDAC1-3 inhibitors, such as CXD101, synergize with anti-PD-1 antibodies to induce an almost complete regression of orthotopic Hepa 1-6 tumors in mice. This antitumor effect may be mediated by the HDAC1-3-induced reduction in MDSCs and enhanced NK cell activation. Further studies are required to clarify the role of MDSCs in enhancing the effects of anti-PD-1 antibodies. These findings reveal a novel mechanism underlying HDAC inhibitors in HCC and suggest a promising combination strategy using anti-PD-1 antibodies for clinical translation.

## Author Contributions

Z.X. conceived and performed the experiments and drafted the manuscript. D. O. contributed to RNA-seq and ATAC-seq experiments. Y.O. and N.O. helped with the experimental design and participated in discussions. M.T. oversaw the study, provided guidance, and contributed to the manuscript writing. All the authors reviewed and revised the manuscript.

## Data Availability Statement

NGS data are available in GEO under the accession number GSE232268. All data are available from the corresponding authors upon request.

