## Supplemental_Data for "HDAC1-3 inhibition reduces CCR2 expression and immunosuppressive function of myeloid-derived suppressor cells"

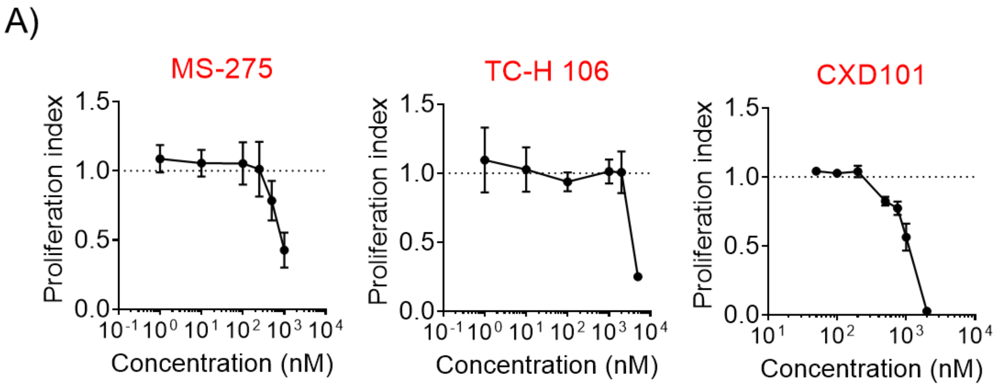


**Fig. S1 The proliferation of MDSCs following with HDAC inhibitor treatment. A)** After 4 days of culture in media supplemented with GM-CSF (40 ng/mL) with or without the addition of different concentrations of HDAC inhibitors, the number of MDSCs was determined using flow cytometry with precision count beads. The proliferation index was calculated as a ratio of the number of MDSCs treated with HDAC inhibitor to that in non-treated group. Data represent mean ± S.E.M., pooled from three independent experiments.

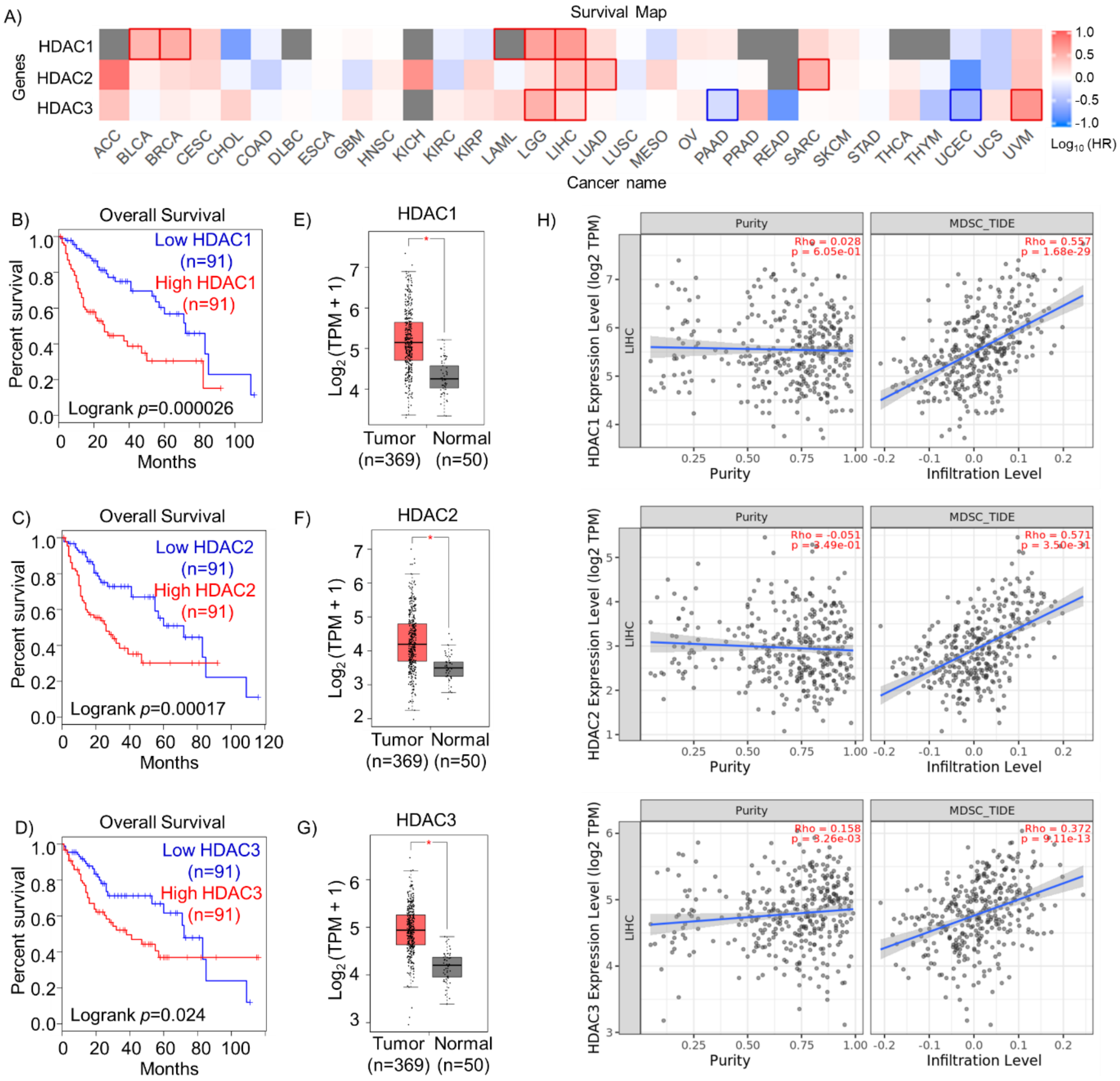


**Fig. S2 High expression HDAC1, 2, 3 are related with poor prognosis of HCC. A)** The survival contribution of *Hdac1*, *Hdac2*, and *Hdac3* in multiple cancer types was assessed by the GEPIA2 web server. The survival heat map shows the hazard ratios (HR) in logarithmic scale (log10) for different genes. The red and blue blocks indicate higher and lower risks, respectively. The rectangles with frames indicate the significant results in prognostic analyses (quartile cut-off, $p < 0.05$ using the Mantel–Cox test). **B-D)** The overall survival curves comparing the high (red line) and low (blue line) (quartile cut-off) expressions of *Hdac1*, *Hdac2*, and *Hdac3* in liver hepatocellular carcinoma (LIHC) in the GEPIA2 web server. **E-G)** The expression of *Hdac1*, *Hdac2*, and *Hdac3* in the LIHC tissue group compared with the normal tissue group. Asterisk represents fold change > 1.5 with $p < 0.01$. The dots represent expression in each sample. **H)** Correlation of *Hdac1*, *Hdac2*, and *Hdac3* expression with tumor purity (the proportion of cancer cells in a sample; left) and with the infiltration level of MDSCs (right) analyzed by the TIMER2.0 web server in LIHC.

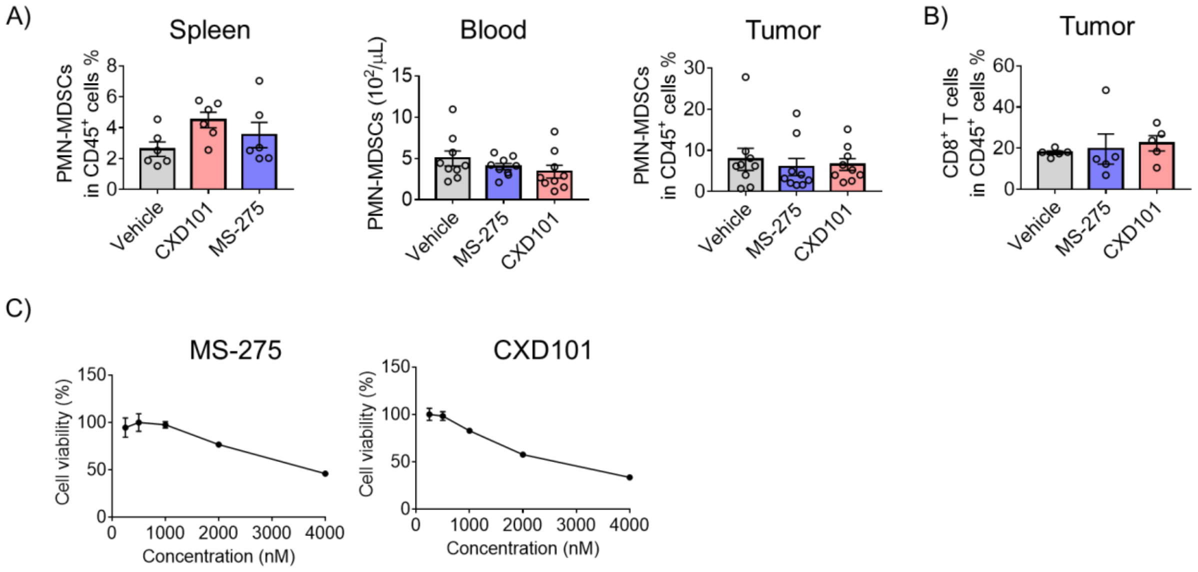


**Fig. S3 A)** Flow cytometry of the PMN-MDSCs ($CD11b^+Ly\text{-}6G^+Ly\text{-}6C^{int}$) in spleen, blood, and tumors. Data represent mean ± S.E.M. of one (spleen) or two (blood, tumor) independent experiments (**p* < 0.05 by one-way ANOVA). **B)** Flow cytometry of the proportion of tumor $CD8^+$ T cells ($CD3\varepsilon^+NK1.1^-CD8\alpha^+$) in $CD45^+$ live cells. Data represent mean ± S.E.M. of two independent experiments (**p* < 0.05, ***p* < 0.01 by one-way ANOVA). **C)** Hepa 1-6 cells treated with MS-275 or CXD101 for 48 h at various concentrations *in vitro* and cell viability was then measured using the CCK-8 kit.
